# Does force-field adaptation induce after-effects on space representation?

**DOI:** 10.1101/238246

**Authors:** Carine Michel, Lucie Bonnetain, Olivier White

## Abstract

Prism adaptation is a well-known model to study sensorimotor adaptive processes. It has been shown that following prism exposure, after-effects are not only restricted to the sensorimotor level but extend as well into spatial cognition. The main purpose of the present study was to investigate in healthy individuals whether expansion to spatial cognition is restricted to adaptive processes peculiar to prism adaptation or whether it occurs as well following other forms of adaptive process such as adaptation to a novel dynamic environment during pointing movements. Representational after-effects were assessed by the perceptual line bisection task before and after adaptation to a leftward or a rightward force field. The main results showed that adaptation developed at sensorimotor level but did not produce after-effects in space representation. However appropriate analysis showed that the slower a participant de-adapt to a rightward dynamic perturbation, the stronger the influence on the perceptual midline judgment during the late phase of the bisection task. The discussion highlights the commonalities between prism and dynamic adaptation on the effects on space representation.

## Introduction

Graceful motor actions rely on successful interaction between our body and the environment. When one moves an object, the brain takes into account the physics of the task to adjust the motor commands sent to the arm and the hand. Handing a book to someone is a smooth and effortlessly process. In unfamiliar circumstances, however, early movements can exhibit large errors, because the consequences are very different from what was initially planned. In the latter case, this discrepancy - n error signal - is used by the brain to drive adaptation such that smooth movements are eventually restored. This generic mechanism has been generalized to many types of actions ^1-3^.

One of the classical and oldest models to study sensorimotor adaptive processes is prism d ptati’on. Participants wear prisms that deviate the visual field laterally while they point to visual targets ^4^. Initially, subjects make pointing errors in the direction of the optical deviation. On the basis of these error signals, subjects gradually improve their performance until they achieve an accurate behavior. When the prisms are removed, the sensorimotor correlations revert to an inappropriate state and the pointing movements are shifted in the direction opposite to the prismatic shift. The sensorimotor after-effects can be explained by proprioceptive, visual and motor control changes ^5^.

The interest taken in prism adaptation was considerably increased since the demonstration that after-effects are not restricted to the sensorimotor level but that they extend as well into spatial cognition. The term ‘cognition’ is not picked up randomly: it refers to the fact effects are not bound to the usual framework of compensatory sensorimotor after-effects but also involve mental abilities such as judgement, comparison or mental representation of space. It is worth underlying that in healthy individuals, cognitive after-effects occur following adaptation to a leftward optical deviation. For example only adaptation to leftward optical deviation turns representational pseudoneglect (leftward bias in midline judgments corresponding to a mental over-representation of the left part of space) into neglect-like behavior (rightward bias in midline judgments corresponding to a mental over-representation of the right part of space). Neglect simulation was not only described in peripersonal (i.e. within arm-reach, ^6^,^7^), extrapersonal (i.e. beyond arm-reach, ^8^,^9^) and bodily space representation ^10^ but also in mental numbers ^11^ and letters scales ^12^. The influence of prism adaptation extends also to spatial attention ^13^, hierarchical processing ^14^,^15^ and spatial remapping ^16^,^17^.

Here, we set out to investigate whether the asymmetrical expansion of sensorimotor after-effects induced by prisms to spatial cognition is a phenomenon specific to prism adaptation or whether it occurs as well following other forms of adaptive processes such as when we adapt pointing movements to a novel dynamic environment. If neglect-like cognitive after-effects in healthy individuals depend on the occurrence of sensorimotor effects, then, we should observe spacerepresentational biases only after dynamic adaptation to a leftward force field that requires a compensatory rightward motor command. In contrast, if the generalization of adaptation to spatial cognition depends on adaptive processes peculiar to prism adaptation, after-effects of adaptation to force field should not interfere with space representation.

## Results

In the present study we investigated both sensorimotor and cognitive after-effects following adaptation to velocity-dependent forces perpendicular to vertical arm movements. Cognitive after-effects on spatial representation were assessed with the classical perceptual line bisection task ^18^,^19^.

### Sensorimotor adaptation to force field

Participants reached toward 3 targets vertically aligned but displayed at three distances from the start position. We formed two groups (see Fig. 1, G_Left_ and G_Right_) depending on the direction of force field perturbation (Left vs. Right). Overall, participants achieved peak velocities in line with instructions (mean±SD, 70.7±2.8, 78.9±4.2 and 89.4±6.9cm/s for the near, medium and far targets, respectively). During the training session (Fig. 1, “Training” and Fig. 2A) and for each group separately, we did not observe significant effects of trials or targets on angular errors 150ms after movement onset (Trials: G_Left_ F_53_,_597_=1.0, p=0.469; G_Righ_, F_53_,_597_=0.8, p=0.796 and Targets: G_Left_, F_2_,_45_=0.1, p=0.871; G_Right_, F_2,42_=0.1, p=0.890). A direct comparison between these two groups yielded no difference either (independent t- test, t_29_=-1.1, p=0.151).

**Figure 1:**
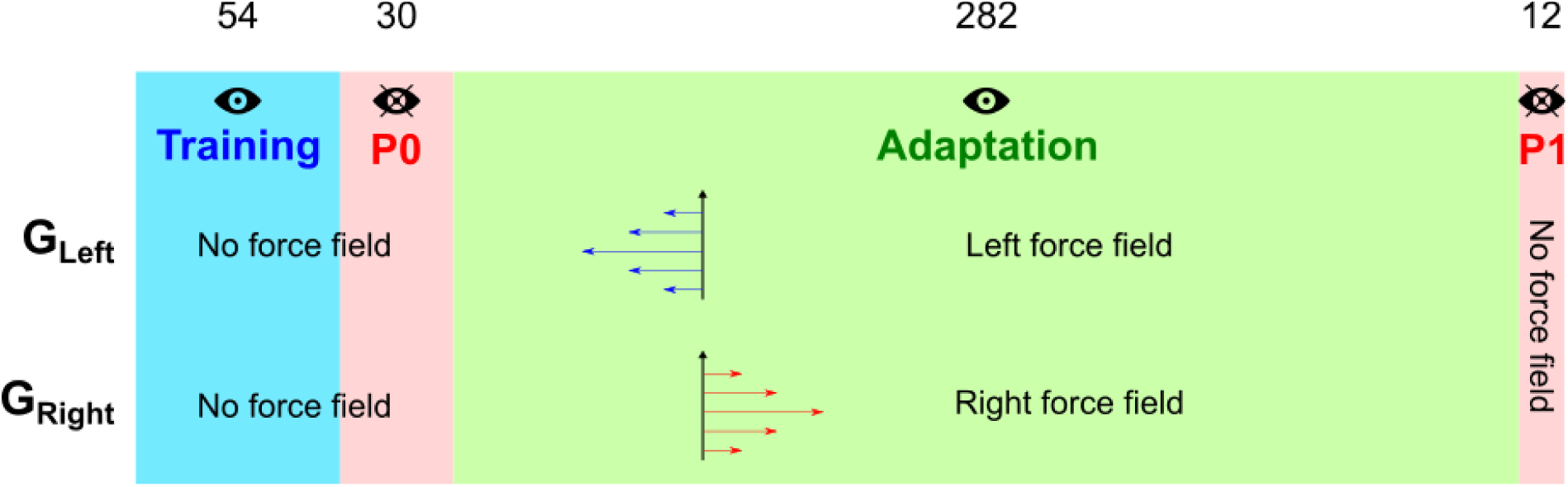
Experimental procedures. The four rectangles correspond to the four experimental sequences. Number of trials is reported above each sequence. Each group (G_Left_ and G_Right_) were subjected to different force field perturbation directions as depicted by the blue and red horizontal arrows in the “Adaptation” sequence. An “eye” icon corresponds to the presence of visual feedback information about the cursor trajectory. “PO” and “PI” stand for “Probe before adaptation” and “Probe after d ptati’on”, respectively.

**Figure 2:**
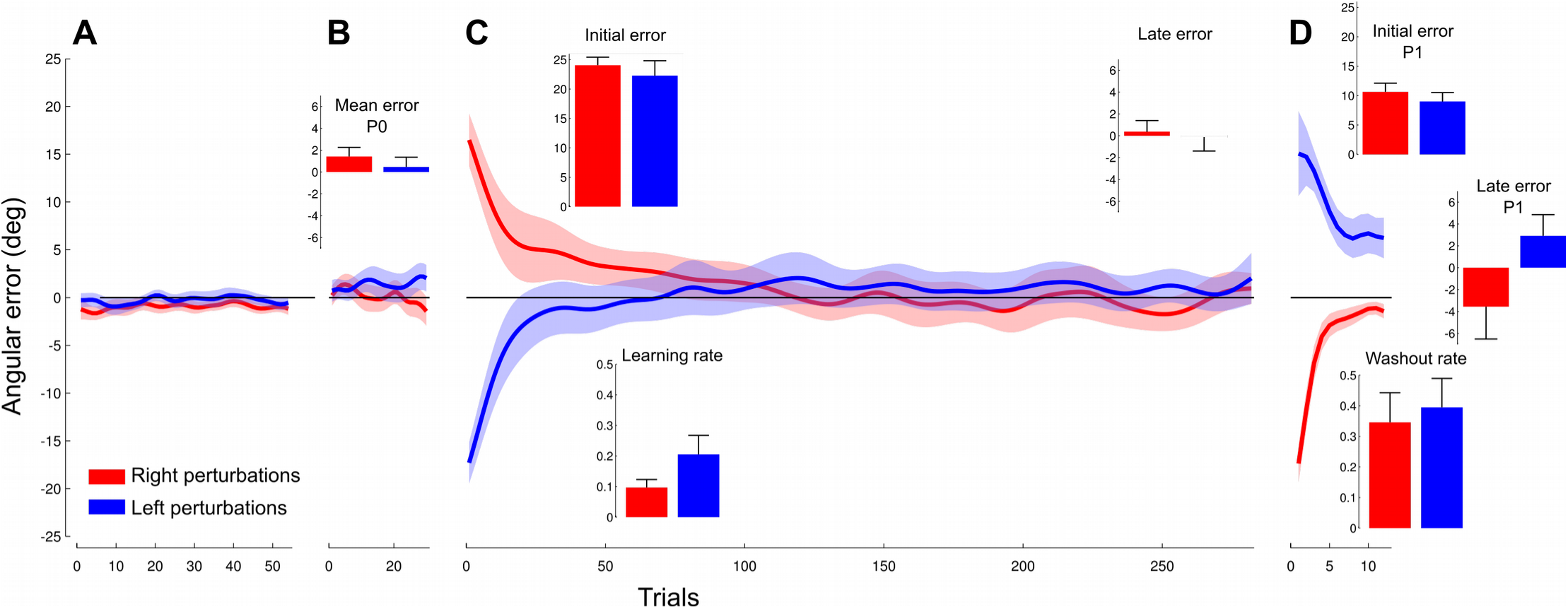
Angular errors 150ms after movement onset during the four phases of the “Adaption” session for the group who experienced rightward (red) or leftward perturbation (blue). (A) Baseline trials show that errors fluctuate around zero. (B) First open-loop phase. The inset presents the average of the errors for the block. (C) Errors decrease from positive (right group, red) or negative (left group, blue) values to close to zero. Initial error and late error insets report the average initial and final errors, respectively. Learning rates calculated for each subject on the exponential fit are shown below the x- axis in panel C. (D) Second open-loop phase. Initial errors highlight after-effects in the opposite direction than initial errors in the “Adaptation”. Note that late errors did not vanish completely. Washout rates are also reported for the sake of comparability. Error bars and shaded areas correspond to SE.

Participants then performed 30 trials without any visual feedback about trajectory (Fig. 1 “PO” and Fig. 2B). This short block allowed us to assess open loop performance. Again, we failed to find any difference across trials (G_Left_, F_29,362_=0.5, p=0.978; G_Right_, F_29_,_344_=0.9, p=0.642) or targets (G_Left_, F_2_,_45_=0.4, p=0.700; G_Righ_t, F_2,42_=0.01, p=0.991) on the initial angular error and both groups were not different (independent t-test, t_29_=−0.8, p=0.227).

We then exposed participants to a large block (282 trials) during which they underwent a leftward (G_Left_) or rightward (G_Right_) velocity-dependent force field that perturbed movement trajectory (Fig. 1 “Adaptation” and Fig. 2C). Initially and expectedly, both groups made large errors in the direction of the perturbation (first three trials, G_Left_: −24.1±4.9deg; G_Right_: 22.3±8.4 deg) that vanished after a brief exposure period (average for the last 30 trials, G_Le_ft: −0.1±5.0deg; G_Right_: 0.4±4.2 deg). Both groups exhibited the same behavior in terms of (unsigned) initial and (signed) final absolute errors (independent t-test, initial error t_20_=−0.6, p=0.273, final: t_28_=−0.3, p=0.385) and adaptation rates (mean=0.16; t_29_=1.5, p=0.070).

Participants ended this session with a second 12-trial block without any visual feedback about performance (Fig. 1 “PI” and Fig. 2D). Since the perturbation was absent in that last block, the initial errors reflected feedforward behavior. Both groups made large errors in the opposite direction of the perturbation (first three trials, G_Left_: 10.6±5.7deg; G_Right_: −9±5.7 deg) that decreased over trials. Groups had similar performances for initial and final errors (independent t-test, t<0.9, p>0.178) and washout rate (mean=0.37, t_29_=0.3, p=0.367). Importantly, we found significantly although borderline larger errors at the end of the PI phase compared to baseline errors in the PO phase in the G_Left_ (paired t- test, t_14_=1.8, p=0.049,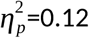) and G_Right_ groups (paired t-test, t_15_=1.8, p=0.042, 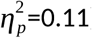). In other words, participants of either group were not fully de-adapted at completion of the washout phase and hence before entering the second bisection test.

### Baseline space representation assessed by the first bisection task

Participants always provided judgments of the location of the tick within 3000ms after stimulus presentation. However, this timing was influenced by two experimental factors. On the one hand, response time was modulated by tick offset following a bell-shaped curve. Timing increased from low values (∼1300ms) for large offsets to much longer latencies (∼2600ms) when the offset was nearer midline and therefore harder to judge. On the other hand, participants were on average 17% faster to react during the second session (Bisection-POST). A 2-way repeated measure ANOVA confirmed these main effects (Offset: F_16,1o2o_= 16.7, p<0.001,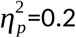; Session: F_16,102o_=32.3, p<0.001,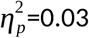) without interaction (F_16_,_102_o=0.5, p=0.953).

We estimated, for each participant, the threshold that corresponded to the offset for which the probability of correct response was 50% (see Methods). We did not find any difference between groups (independent t-test, t_29_=-1.2, p=0.249). Furthermore, the thresholds in this initial BISECTION-PRE session were different from an offset of 0mm (mean=−0.08mm, SD=0.29mm, independent t-test: t_60_=- 2, p=0.025,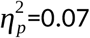). Both groups had comparable performance before entering the ADAPTATION session and estimated mid-segment slightly on the left.

### Effects of sensorimotor adaptation on space representation

We now ask the question as to whether the point of subjective equality was influenced by sensorimotor adaptation induced by the force field. We failed to report any difference between thresholds before and after the adaptation phase (BISECTION-PRE VS. BISECTION-POST, F_144_=0.5, p=0.598) or between groups (G_Left_ vs. G_Right_, F_144_=0, p=0.982). Furthermore, the ANOVA did not report any interaction between group and the bisection task on these thresholds (F_144_=0, p=0.958).

We adopted a more sensitive approach to try and detect any difference between these thresholds. For every participant, we fitted a sigmoid function and regressed the offset that corresponded to chance level to measure the subjective estimation of the line center. Figure 3A depicts the fit for all participants together. Note that the red and blue cursors are superimposed and correspond to thresholds for BISECTION-PRE (blue) and BISECTION-POST (red), respectively, calculated on the whole dataset. We defined a new statistics as the difference between the thresholds calculated during BISECTION-PRE and BISECTION-POST phases and we averaged these values across participants. We then ran a bootstrap procedure (50,000 repetitions) to approximate the population (Fig. 3B). We could not reject the null hypothesis (no difference between the means, Fig. 3B, blue cursor). Indeed, the true mean (Fig. 3B, yellow cursor, 0.01) is well contained within the 95%-confidence interval (Fig. 3B, red cursors, [−0.13 to 0.15], p=0.895). In sum, these analyses show that a sensorimotor adaptation to a force-field did not influence the perceptual thresholds in the line bisection task.

**Figure 3:**
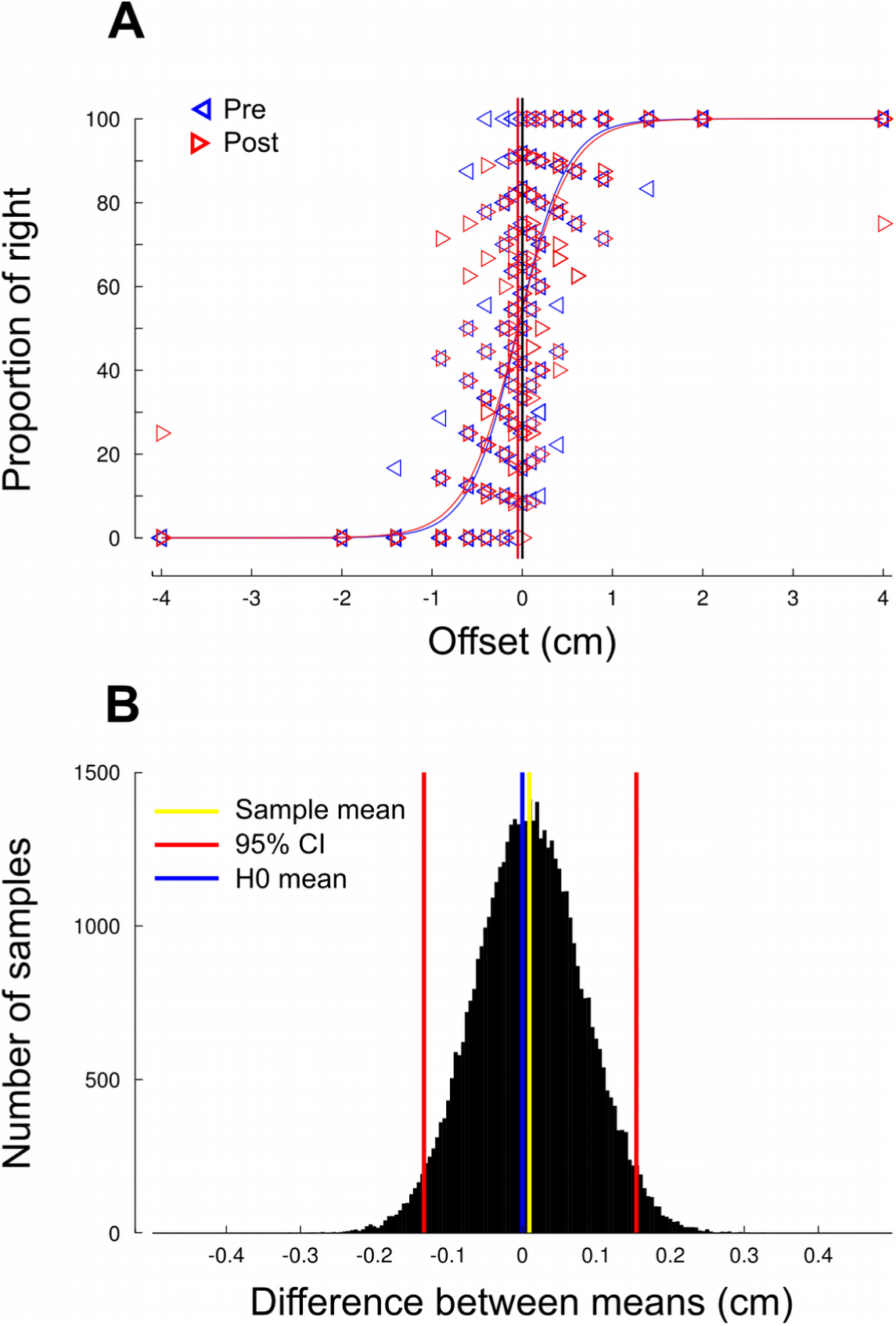
No difference between bisection thresholds before and after force field adaptation. (A) Sigmoidal fit for all participants, before (Pre, left blue triangles) and after adaptation (Post, red right triangles). (B) Bootstrapped population (50,000 repetitions) of the difference between thresholds before and after adptation. The estimated population mean (yellow cursor) is not different from 0.

### Perceptual thresholds and washout rates

Why did we fail to highlight an effect of a strong force field adaptation task on the perceptual thresholds? One reason may be because participants partially de-adapted during the Pl-phase. Indeed, there were reminiscent errors at the end of PI compared to P0, but they just hit statistical significance. In other words, only a few trials without perturbation and without visual feedback of the performance were sufficient to significantly decrease the error (see Fig. 2D).

However, all participants did not de-adapt the same way. We pushed our investigations one step further and coupled data from the three experimental sessions. We reasoned that the slower a participant de-adapt, the stronger the influence on her/his perceptual judgment on the second bisection task. To quantify this, we correlated the absolute difference between thresholds after and before the adaptation session (|T_p0st_-T_pre_|) and the washout rate (parameter a_2_ calculated in the PI phase). The spearman correlation yielded no significant relationship between a shift in threshold and the washout rate (Fig. 4A, r=0.15, p=0.450). However similar in many ways, the two groups were exposed to left and right dynamic perturbations. The same correlation calculated on each group separately provided no significant outcome either (G_Left_: r=0.3, p=0.923; G_Right_: r=0.31, p=0.254). Therefore, we cannot conclude that a reminiscent adapted state led to a shift in thresholds nor that there was an influence of direction of perturbation on these perceptual thresholds.

**Figure 4:**
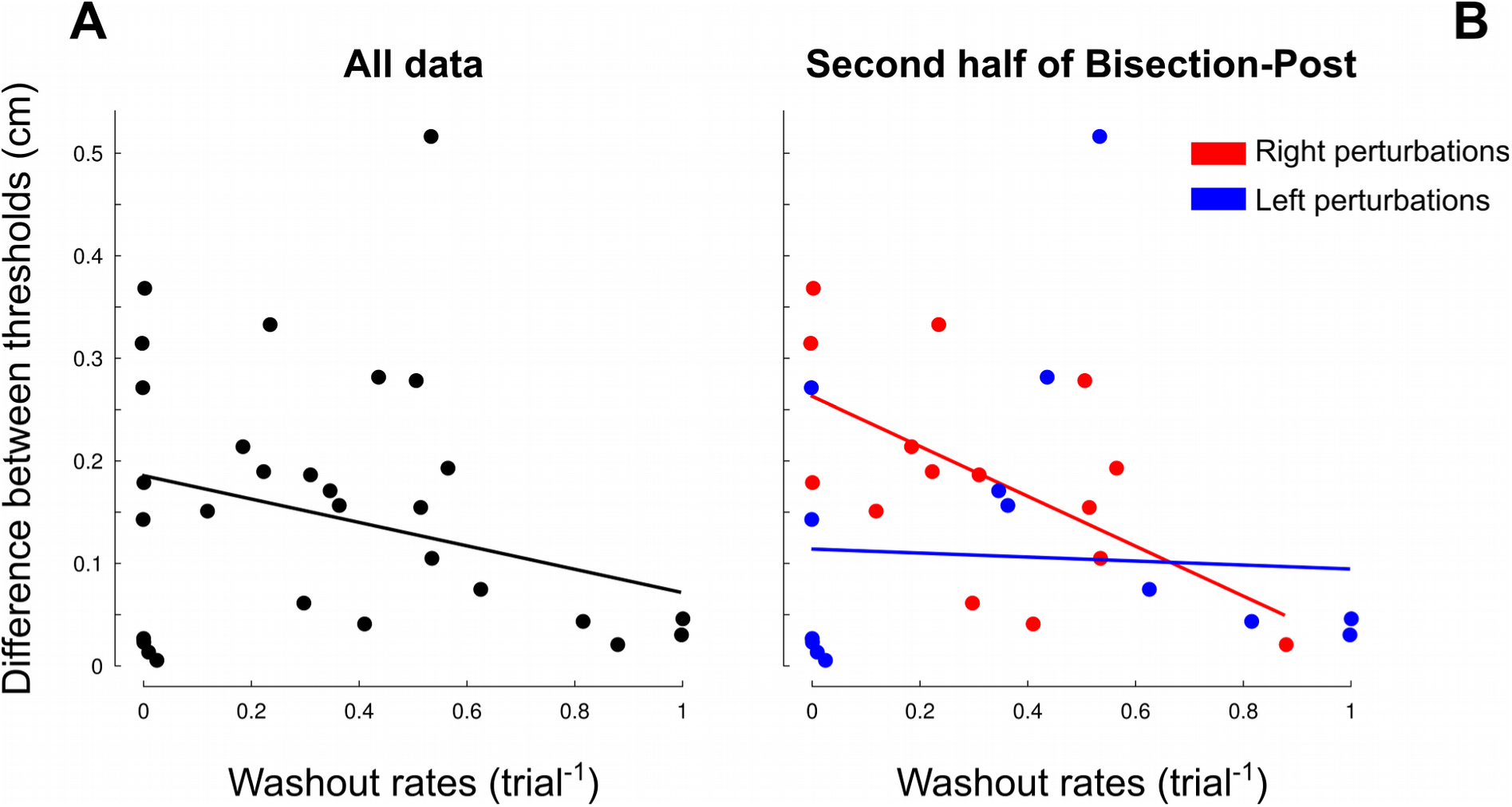
Spearman correlation between washout rates (x-axis) and absolute difference of estimated thresholds between the two bisection sessions. (A) No correlation when the whole dataset is considered. (B) Correlation between the same variables but only for the second half of the BISECTION-POST session and separated by group (see legend). The slope and offset of the linear model were −0.24 and 0.26 for the right group (red) −0.02 and 0.11 for the left (blue) group.

The passage of time is known to reinforce internal motor memories ^20^ and after-effects have been shown to emerge some time after being exposed to perturbations ^21^. We conducted the same n lysis but we considered separately the first half (68 trials) and the second half of trials during BISECTION-POST. We experimentally designed the BISECTION-POST block in such a way to ensure balanced distributions of offsets. Quite surprisingly, whereas we did not find a significant correlation during the early BISECTION-POST phase (G_Left_: r=0.47, p=0.088; G_Right_: r=0.09, p=0.754), we found an effect in the late BISECTION-POST phase. Indeed, only the washout rates of the group that was perturbed to the right - and had to compensate the force field by pushing to the left - were correlated with the difference in thresholds (Fig. 4B, G_Left_: r=0.09, p=0.771; G_Right_: r=0.53, p=0.036). In other words, Figure 4B shows that the larger a_2_ (or the faster the washout), the smaller the difference between perceptual thresholds.

## Discussion and conclusion

The main objective of the present experiment was to investigate whether adaptation to a dynamic perturbation transfers to space representation in healthy individuals. We failed to show that d ptati’on developed at sensorimotor level influenced after-effects in space representation. However, we found that the slower a participant de-adapt to a rightward - and not leftward - force field perturbation, the stronger the influence on the perceptual judgment on the late phase - and not early phase - during the bisection task.

### Replication of sensorimotor adaptation and presence of baseline pseudoneglect

Robotic devices are commonly used to highlight sensorimotor adaptation by perturbing motor tasks using different forms of dynamics such as elastic ^22^,^23^, viscous ^3^, inertial^24^ or composite force fields ^25^. Here, participants exhibited the classical adaptive behaviour to novel dynamic environments: large initial errors in the direction of the perturbation and then large errors in the opposite direction of the expected perturbation, when it is removed unbeknownst to the participants. Here, we set high viscous gains and asked participants to perform reaching movements of large amplitudes. The combined effects of high velocity toward the target and large viscous gain put us in a position to amplify adaptation. Even if the amplitude of after-effects decreased over trials in the open-loop reaching following adaptation (“PI” period), participants were still partially adapted at completion of the washout phase. Participants were therefore in a state in which their internal model was still partially adjusted to the task dynamics. This experimental paradigm leaves us on sound ground to test whether this dynamic sensorimotor adaptation interferes with space representation.

The occurrence of cognitive after-effects in healthy individuals seems to depend on the baseline expression of pseudoneglect ^26^,^27^. We took the precaution to assess pseudoneglect by means of the perceptual judgment version of the line bisection task (landmark test) to prevent any active or passive movement of the participant. Furthermore we used long segments (400mm) because they can capture subtle representational effects ^*2B*^. In agreement with previous studies, we also report that participants exhibited a leftward pseudoneglect bias due to a mental over-representation of the left part of space ^*2S*^. However, its effect size was smaller compared to other studies ^21^. One reason may be because we did not impose a stringent time constraint to provide the judgment, leaving more time to the participant to make a decision. The experimental conditions are therefore optimal to assess if a strong sensorimotor adaptation to force fields influences space representation.

### Sensorimotor adaptation and space representation: two independent processes?

In spite of both favourable sensorimotor and cognitive experimental conditions, we failed to observe obvious cognitive after-effects following adaptation to a strong dynamic perturbation. Sensorimotor d ptati’on to both leftward and rightward deviating prisms share a similar effect as force field d ptati’on in that significant sensorimotor after-effects are observed once the perturbation is removed. However, a peculiar difference resides in the fact that prism adaptation - when the optical deviation is at least lOdeg ^29^ - transfers to more cognitive dimensions such as mental scales or spatial attention and untrained sensory domains (see Introduction). Furthermore, this effect is asymmetrical: only leftward deviating prisms induce a rightward shift in the perception of segment midline ^6^,^10^,^13^,^26^. What fundamental differences between these two types of adaptation are responsible for this lack of cognitive effect?

For years, prism adaptation and the reason why after-effects are observed have been described through a dual-process framework. Initially, a rapid process occurs (referred to as ‘calibration’) and allows to reduce the error by using feedback (proprioceptive and/orvisual) to better characterize what is going on. This process is responsible for a quick decline of the initial error. In parallel, a slow process (referred to as ‘realignment’) brings back the different reference frames in congruence. Indeed, under prismatic shift, the visual/motor reference frame is affected while the proprioceptive/motor reference frame is not. The first process is cognitively demanding and contributes little to prism after-effects. The second, more automatic, process develops gradually and is thought to be mainly responsible for prism after-effects. An inescapable experimental support to the distinction between fast and slow processes was provided by a detailed analysis of arm movement trajectories ^30^. The two compensatory processes could be distinguished within a single reaching movement. The fast strategic compensation of the optical shift consists of trial-by-trial updating of the planned movement direction on the basis of the error experienced during the previous trial. This compensation is a strategic demanding process that drives error correction early during prism exposure and does not generate after-effects. In contrast, the transformation of spatial maps (visual, proprioceptive and motor) to bring origins of coordinate systems into correspondence is n utomati’c process that develops more gradually during prism exposure and correlates with after-effects measured at the end of the experiment. These two processes operate relatively independently from one another (O’Shea et al., 2017).

More recently though, this two-ti’mescale process was challenged. Indeed, a third ultra-slow process was needed to explain the immediate and long-term retention of prism after-effects when prismatic exposure reached 500 trials ^31^. More generally, there may be a continuum of learning processes that are combined according to a number of contextual factors. In support of this view, differentti’mescales in neural network dynamics have been shown in typical adaptation paradigms ^32^,^33^ and models ^34^. Furthermore, in other motor adaptation paradigms (visuomotor rotations or force field d ptati’on), the retention of after-effects also depends on the experimental context, such as if we learn using error-based mechanisms or use-dependent mechanisms ^35^.

Even if none of these processes explain why prism adaptation generates cognitive after-effects while force field adaptation does not, we may hypothesise that the main difference between prism and force field adaptation may lie in the relative contribution of the fast and slow processes. Indeed, it has been shown that the occurrence of cognitive after-effects depends on the development of the slow process ^7^. In prism adaptation, the slow process may dominate and be responsible for strong, persistent and generalized sensorimotor and cognitive after-effects ^17^. In force field d ptati’on, the slow process may be minor and responsible for short-lasting and poorly generalized sensorimotor after-effects and non-significant cognitive after-effects. Indeed, adaptation to force field is not only spatially highly contextual but is also restricted to the exposure during training ^36-38^.

From a neurophysiological point of view, common neural substrate for both adaptation to force field and to prismatic shifts involved neural structures well-known to be implicated in sensorimotor processes such as the cerebellum and the motor cortex ^39-41^. Nevertheless, the impressive generalization of prism adaptation to high-order processes are mediated through the modulation of cerebral areas involved in spatial cognition via a bottom-up cerebello-corti’cal network involving temporal cortex ^42^ or posterior parietal cortex (inferior parietal lobule) ^21^,^43^,^44^. it is likely that the slow process developed during dynamic perturbation is not strong enough to mobilize such a bottom-up cerebello-corti’cal network underpinning cognitive processes.

### Delayed effects of right force field perturbations on space representation

Although we did not observe, on average, an effect of sensorimotor after-effects on the perceptual task, we identified a specific condition in which a shift between midline judgment thresholds were present. Indeed, we found that the more participants are in an adapted state (small washout rate), the more the thresholds are shifted to the right. This finding however only held for rightward perturbation and in the second part of the bisection task.

Prism adaptation induce asymmetrical effects on space representation as demonstrated by rightward deviations after being exposed to leftward prismatic shifts. Here, we also found an asymmetrical effect of force field adaptation to space representation. What processes shared between prism and force field adaptation could explain this result? Under leftward visual shifts, the hand must point to the right of the visual target to hit it. The reach trajectory remains identical in space and the target becomes only proprioceptive. When subject to rightward force field, the motor command must be adapted so as to counteract the expected perturbation. Therefore, the brain adds a compensatory force profile to the left, on top of the baseline motor command. The target position is unchanged both in the visual and proprioceptive spaces. Like in prism adaptation, the reach trajectory in space is undisti’nguishable from baseline movements. To sum up, after adaptation, the outcome movement in space is the same in both cases. However, the strategic and motor compensations to generate that movement rely on different visual and proprioceptive information in the two approaches.

Bayesian theory tells us how to combine uncertain sources of information to plan the most accurate action. For instance, one can rely on vision and proprioception. However, the reliability of these two modalities can be different^45^. Specifically, the brain weights each source of information by the inverse of their uncertainty. Each modality has a prior (prediction) and a likelihood (observation) that are then combined in a multimodal integrator to yield the most likely estimate of the hand location. In the first stage of prism exposure, the visual hand and target estimates are shifted but the proprioceptive information is unaffected. However, in force field adaptation, visual feedback remains intact - there is no optical shift - while proprioception is perturbed. Therefore, in both cases, some time is necessary to solve a sensorimotor conflict. In prism adaptation, participants dedicate more cognitive resources to the left hemi space, where the left visual target remains visible, and is used to estimate the position of the real target. Asymmetrical neural circuitry explains why this does not hold in rightward prismatic shifts ^46^. We suggest that similar asymmetric, although less powerful, mechanisms occur in force field adaptation. Indeed, cognitive resources may be mobilized first to understanding the effect of the rightward perturbation to process uncertain proprioceptive information since the cause of the disturbance is clearly attributed to proprioception and not vision ^47^,^48^. Then, the more participants adapt, the more refined become their leftward compensatory motor command. Altogether, we suggest that in both cases and in the adapted state, cognitive attention between the two paradigms is biased to the left hemi space. In prism adaptation, participants monitor the visual distance between the proprioceptive and (left) visual targets. In force field adaptation, participants program a strong leftward motor command. We speculate the asymmetry observed in the second case share peculiarities as prism adaptation. Furthermore, the weaker effect of the asymmetry in the dynamic case could be explained by the different modalities on which the brain mustfocus to maintain the adaptation state.

Finally, we also found that these effects emerged sometime after adaptation. Whereas most studies report immediate after-effects ^6^,^10^, one found that space representation after-effects developed after 5 minutes and lasted for up to 40 minutes ^21^. A more recent study showed representational after-effects that lasted until 8 hours (Schintu et al., 2017). Here, the first trial in the second block occurred 6 minutes after the end of the adaptation phase, which is a delay comparable to the one observed above. This process may reflect the time needed by cortical networks to make links with the areas involved in space representation. This is also in line with the intermittency observed by Schintu and colleagues in the after-effects on space representation since specific areas, differentfrom the ones involved in motor adaptation, are recruited.

Here we show that force field adaptation shares a similar mechanism as prism adaptation in that it induces asymmetrical shift in midline judgment that emerge a few minutes after reaching the end of the adapted state. The asymmetrical effects were however small for two possible reasons. First, we tested participants in the second bisection task after partial de-adaptation. Therefore, we predict magnified effects if the quality and robustness of the adapted state are optimized. This could be achieved by inducing gradual dynamic perturbations for a larger number of trials and by testing participants immediately without washout phase. Second, the weaker asymmetry may be caused by the fact a focus on visual information (prism adaptation) induces stronger biases than a focus on the motor command (dynamic perturbation). Follow-up experiments, testing participants under optimized adapted state and using visuomotor adaptation instead of force-field adaptation should address these issues

## Materials and Methods

### Participants

Thirty-one self-reported right-handed adults (16 females, 15 males, mean age=24.3, SD=4.9 years) participated voluntarily in the experiment. All subjects were healthy, without neuromuscular disease and with normal or corrected to normal vision. The experimental protocol was carried out in accordance with the Declaration of Helsinki (1964) and the procedures were approved by the local ethics committee of Université de Bourgogne-Franche Comté. All participants were naïve as to the purpose of the experiments, were debriefed after the experimental sessions and received a USB stick for their participation.

### Experimental procedure and apparatus

The experiment consisted in three separate phases (BISECTION-PRE, ADAPTATION AND BISECTION-POST), described below.

During the first, BISECTION-PRE phase, participants faced a high resolution LCD screen (27” 1440×2560) in a dimly illuminated room. Participants were asked to perform a bisection segment test ^19^. A total of 130 horizontal green segments (length: 400mm, thickness: 2mm) were sequentially displayed on the screen together with a perpendicular green tick (height: 30mm, thickness: 2mm). Participants were requested to judge (forced-choice) whether pre-transected lines were bisected to the left or to the right of their centre. The response was given verbally. The experimenter manually recorded responses by hitt’ng the appropriate key on a standard computer keyboard (‘L’ for left and ‘R’ for right) to limit any motor action for the participant. Reaction time was recorded as the time between stimulus onset and key strike by the experimenter. No response provided before a 20-s timeout was considered as missing information. Ticks offsets were not uniformly distributed but followed a Gaussian shape in order to increase sampling around the Euclidian center and, hence, enhance sensitivity of subsequent data analysis (Table 1).

**Table 1:** Distribution of perpendicular ticks offsets with respect to the Euclidian center of the segment (Omm). The first bold row reports the offsets (mm). The second row (BISECTION-PRE) reports the number of times a tick was presented with that offset in the 130-trial block. The last two identical rows present the offset distribution during the first (BISECTION-POST, 1^st^ half) and second half (BISECTION- POST, 2^nd^ half) of the 136-trial block. The distribution was Gaussian and centered on the veridical line center. Note that the largest offsets (±40mm) corresponded to 10% of the length of the horizontal segment.

|  | Offset (mm) |  |  |  |  |  |  |  |  |  |  |  |  |  |  |  |  | Total |
| --- | --- | --- | --- | --- | --- | --- | --- | --- | --- | --- | --- | --- | --- | --- | --- | --- | --- | --- |
|  | -40 | -20 | -14 | -9 | -6 | -4 | -2 | -1 | 0 | 1 | 2 | 4 | 6 | 9 | 14 | 20 | 40 |  |
| **BISECTION_PRE** | 4 | 4 | 6 | 7 | 8 | 9 | 10 | 11 | 12 | 11 | 10 | 9 | 8 | 7 | 6 | 4 | 4 | **130** |
| **BISECTION_POST (1st half)** | 2 | 2 | 3 | 4 | 4 | 5 | 5 | 6 | 6 | 6 | 5 | 5 | 4 | 4 | 3 | 2 | 2 | **68** |
| **BISECTION_POST (2nd half)** | 2 | 2 | 3 | 4 | 4 | 5 | 5 | 6 | 6 | 6 | 5 | 5 | 4 | 4 | 3 | 2 | 2 | **68** |

To prevent any cognitive strategy, participants were specifically told that ticks never occurred exactly on the middle of the segment and that their distribution was not systematically symmetric. Ticks were presented randomly and the computer screen went white for 1500ms between each trial to reset the visual display and prevent the participant from using any cue between trials. At completion of this BISECTION-PRE phase, the participant was blindfolded (to avoid any movement involving visuo-manual coordination) and her/his stool was gently rotated 90-deg rightward in order to use a virtual environment. The experimenter assisted the participant in this maneuver to limit unnecessary movements.

During the ADAPTATION phase, participants faced a virtual environment equipped with a haptic device (Phantom 3.0, SensAble Technologies, USA) with the head on a chin rest. Participants looked into two mirrors that were mounted at 90 degrees to each other, such that they viewed one LCD screen with the right eye and one LCD screen with the left eye. This stereo display was calibrated such that the physical location of the robotic arm was consistent with the visual disparity information. Participants performed reaching movements with the right hand from the same starting position toward one of 3 targets situated 22, 27 or 32cm above the starting position. Reaching to multiple targets concurrently allows slowing down learning rates and therefore increasing sensitivity to capture these parameters. Indeed, in a Bayesian framework, learning rates are up/down-regulated as a function of uncertainty in beliefs, which is underlined by the generalization across multiple targets ^49^. Note that unlike in other similar experiments, we used three targets aligned vertically in order to induce only pure horizontal perturbations (see below). Movements were performed in the natural reaching space in an upward direction, involving shoulder and elbow movements, with the elbow pointing downwards. The 3d position of the robot handle was mapped in real time to a grey cursor (diameter: 3mm). A trial started when the cursor was positioned inside a starting green sphere (diameter: 6mm). Then, upon appearance of a white target (diameter: 6mm), the participant performed a reaching movement to the target. The trial ended when the Euclidian distance between final cursor position and the target was below 4mm and the cursor velocity fell below 1.5cm/s for at least 40ms. We instructed and trained participants to reach the target with a peak velocity (cm/s) within 65-75, 75-85 or 85-95 for the near, medium and far targets, respectively. After each trial, the target turned red or blue if movement was too fast or too slow respectively according to the speed criteria. The robot then gently pushed the hand back toward the starting position to prevent participants from planning and performing active movements and therefore, de-adapting.

After briefing, participants were familiarized with a practice block of 54 trials to the three targets. Cursor trajectories were displayed. These data were not included in the analysis. During the experimental session, participants performed 378 trials in the following order (Fig. 1). The training block consisted of 54 reaching movements (18 to each target) to randomly selected targets (Fig. 1, blue “Training” sequence) with full visual feedback of the cursor trajectory. Then, a smaller 30-trial probing block was presented in the same condition except that visual feedback of the cursor trajectory was withdrawn (Fig. 1, pink “PO” sequence). This sequence allowed us to measure initial feedforward performance in the task. A strong velocity-dependent perturbation was then introduced for the next 282 reaching movements (94 trials per target, Fig. 1 green “Adaptation” sequence). The robotic arm generated a force *(F*_x_) in the horizontal direction (perpendicular to the movement trajectory) that was proportional to the forward velocity *(V*_*y*_) of the hand following *F*_*x*_ *= ±bV*_*y*_, with *b — 7 Ns*/*m.* Participants used visual information about the cursor trajectory to compensate this perturbation over trials. Finally, a second probing block including 4 trials to each target (only 12 trials in total) was presented in the same condition as PO (Fig. 1, pink “PI” sequence). This last sequence, without force field perturbation and without visual information, reliably quantified after-effects. We voluntarily limited the length of this last sequence for two reasons. First, we were not interested by the dynamics of the wash-out phase. Second, and more critically, we intended to test our participants in the third part of this experiment while motor planning was still calibrated to counteract the learned perturbation, i.e., before wash-out took place. Indeed, complete de-adaptation occurs usually during the first 20 trials once the perturbation is removed ^35^,^50^.

The 378 trials were equally distributed between 9 blocks of 42 trials to limit fatigue. We designed the blocks such that a change of condition never occurred between blocks but within blocks. The perturbation was a rightward (positive) force field for 16 participants (Fig. 1, G_Right_) and a leftward (negative) force field for 15 participants (Fig. 1, G_Left_). At completion of this second phase, the experimenter assisted the participant to bring her/him back in the position s/he had during the BISECTION-PRE phase. Again, the participant was blindfolded during the 90 deg leftward rotation maneuver.

The apparatus, instructions and task in the last phase (BISECTION-POST) were identical to those used in the BISECTION-PRE phase except for the number of segments presented on the screen. In this third phase, a total of 136 horizontal green segments were displayed on the screen (there were 130 trials during BISECTION-PRE). This intentional 6-trial difference was due to the fact we partitioned the distribution of tick offsets equally between the first and second halves of the block (Table 1). This design allowed to specifically assess representational after-effects in the early period (from trial 1 to trial 68) and late period (from trial 69 to trial 136) following sensorimotor adaptation to force field.

### Data processing and analysis

In the BISECTION-PRE and BISECTION-POST phases, we recorded participants’ verbal responses for every trial (*“Is the tick positioned to the right or left of the veridical segment midline?*”). The time at which the experimenter hit the ‘L’ or ‘R’ stroke allowed us to have an estimate of participant’s reaction times. For each phase, we calculated the proportion of ‘RIGHT’ responses in function of the offset. This S-shape function saturated at 0% for large negative (left) offsets and 100% for large positive offsets (right). Indeed, the further the offsets from the segment midline, the lower the probability to make erroneous judgments.

To quantify the offset that corresponded to chance level (50%), i.e. the subjective perceptual estimation of the line center, we fitted logistic functions separately through BISECTION-PRE, BISECTION- POST, BISECTION-POST 1^st^ half and BISECTION-POST 2^nd^ half data sets, and for each participant separately (r^2^=0.96 on average), 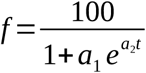 where a_1_and a_2_ were regressed (function nlinfit of Matlab) and t correspond to the offset. The threshold that corresponds to *f* = 50 was calculated according to 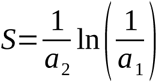 We used two tailed paired t-tests and bootstrapping methods to compare thresholds between BISECTION-PRE and BISECTION-POST and between the two halves ofthe BISECTION-POST phase.

During the ADAPTATION phase, cursor positions were recorded with a sampling rate of 500Hz. Movement start was detected when movement velocity exceeded 3cm/s for at least 100ms. Direction error of each movement was defined as the angle between straight ahead and the segment connecting the start position to the position of the cursor 150ms after movement onset. Such delay reflects mostly the predictive component, i.e. before any feedback could be used to correct the movement^51^,^52^.

We quantified the amount of adaptation by looking at the difference between angular errors in P0 and PI, recorded in the same conditions. We calculated this index for each participant. To quantify adaptation rate, we fitted a power model to the initial angle error, *f* (t) = a_1_(1*—a*_*2*_)^*t*^*+a*_*3*_, and regressed the 3 free parameters a_1_, a_2_ and a_3_. The first, second and third parameters quantify amplitude of adaptation, adaptation rate and final error, respectively.

Sample size was fixed at 15+ participants per group before the start of the experiment. Quantile-quantile plots were used to assess normality of the data before using parametric statistical tests. Variables of interest were submitted to different statistical models (repeated measures ANOVAs) according to the effects analyzed (see Results). Independent t-tests were conducted to compare data between both groups (of unequal size) and paired t-tests were used to compare different conditions within groups. We also report partial eta-squared for significant results to account for effect size. Data processing and statistical analyses were done using Matlab (The Mathworks, Chicago, IL).

## Acknowledgments

The authors declare no conflict of interest. This research was supported by the « Institut National de la Santé et de la Recherche Médicale » (INSERM), the « Conseil Général de Bourgogne » (France) and the « Fonds Européen de Développement Régional » (FEDER).

